# Collective Response of ON and OFF Bipolar Cells to Light Contributes to Improved Visual Acuity

**DOI:** 10.1101/2025.03.03.641188

**Authors:** Anuradha Vinayak Pai, Yogesh A. Kulkarni, Jayesh Bellare

## Abstract

In the visual signal relay, retinal bipolar cells transduce neurotransmitter signals from photoreceptor cells into electrical signals for onward transmission to the brain. The electrical activity of bipolar cells is reflected in the b-wave of the electroretinogram which is majorly contributed by ON bipolar cells leading to the currently accepted belief that light-induced signals travel through these cells only. Here, we probe the role of ON and OFF pathways to generate visual percept under the hypothesis that the ON Bipolar Cell fine image (central illumination) is superimposed onto a coarser image formed due to OFF Bipolar Cell (surround illumination). This improves image sharpness through edge detection by unsharp masking, thereby contributing to improved visual acuity. This new finding could be useful in stimulating the retina through sub-retinal implants to restore vision.

## Introduction

It is widely accepted that visual signals in the retina travel via the Photoreceptor - Bipolar - Ganglion Cell pathway. Electrochemical signals from photoreceptor cells bifurcate into ON and OFF bipolar cells at I plexiform. Of these, the ON Bipolar Cells, which include rod and ON Cone Bipolar Cells (ON CBCs) have the highest density at I plexiform. They depolarize robustly to glutamate reduction and contribute to 80% of b-wave in an electroretinogram (ERG) (Strettoi et al., 2010). Therefore, a reduction in the b-wave of the electroretinogram is interpreted as a loss of vision implying light-induced signals are not propagating.

Besides the ON CBCs, Photoreceptor cells also innervate OFF Bipolar Cells (OFF CBCs / OFF BCs) which contribute to 30% of bipolar cells at I plexiform (Strettoi et. al., 2010). However, the OFF CBC signals have small and negative amplitudes (hyperpolarization), that are generally not seen directly in the b-wave of ERG or measured in other ways because it is dominated by very robust depolarization response from ON CBCs. This has led to neglect of the contribution of the OFF CBCs trivializing the role of these cells in the visual signal pathway.

Therefore, it is generally accepted that (1) the visual signal pathway is from photoreceptors to ON CBCs to ON Ganglion cells and thereon to higher visual centres; and (2) OFF CBCs either do not play a role or their role is considered unknown. This is also reflected in the belief that the b-wave response is insensitive to OFF CBCs. Therefore, a chemical agent (like cis-PDA) that reduces only the OFF CBCs response would not be expected to change the b-wave amplitude, whereas a chemical agent (like L-AP4) that reduces only the ON CBCs would be expected to reduce the b-wave amplitude, implying visual signal transmission across PR-CBC-GC pathway is either reduced or lost (Xu et.al., 2013).

Here, we establish that OFF CBCs give a small but very significant contribution to (electrical) image formation at I plexiform. These cells fine-tune and thus modify electrical images formed by ON CBCs at I plexiform. Thereby contributing to finer details of the image which is very important for object identification. Based on our data, we believe it will be worthwhile to stimulate both ON and OFF CBC types in a proper ratio to restore even a modicum of vision.

## Results

### Effect of ON and OFF CBCs to Light Stimulus

To evaluate the response of both ON and OFF CBCs to light stimuli, we injected rat vitreous with 4 µl 100 mM cis-PDA (cis-2,3-piperidine - dicarboxylic acid), a glutamate analog known to suppress signal transmission from photoreceptors to OFF CBCs (also known as HBCs) and horizontal cells as well as between bipolar cells and third-order neurons (Xu et al., 2003) (Fig.1 Supplementary).

Blocking all post-receptoral pathways other than the ON CBC pathway enabled us to observe the response of the ON CBCs to light stimulus (Slaughter & Miller, 1983) (Fig.1Aii). The ON CBC response was smooth, positively polar, and increased with increasing flash intensity (Fig 1Bii., Fig.1C and Fig. 1Ci.) (Xu et al., 2003).

**Fig. 1.**
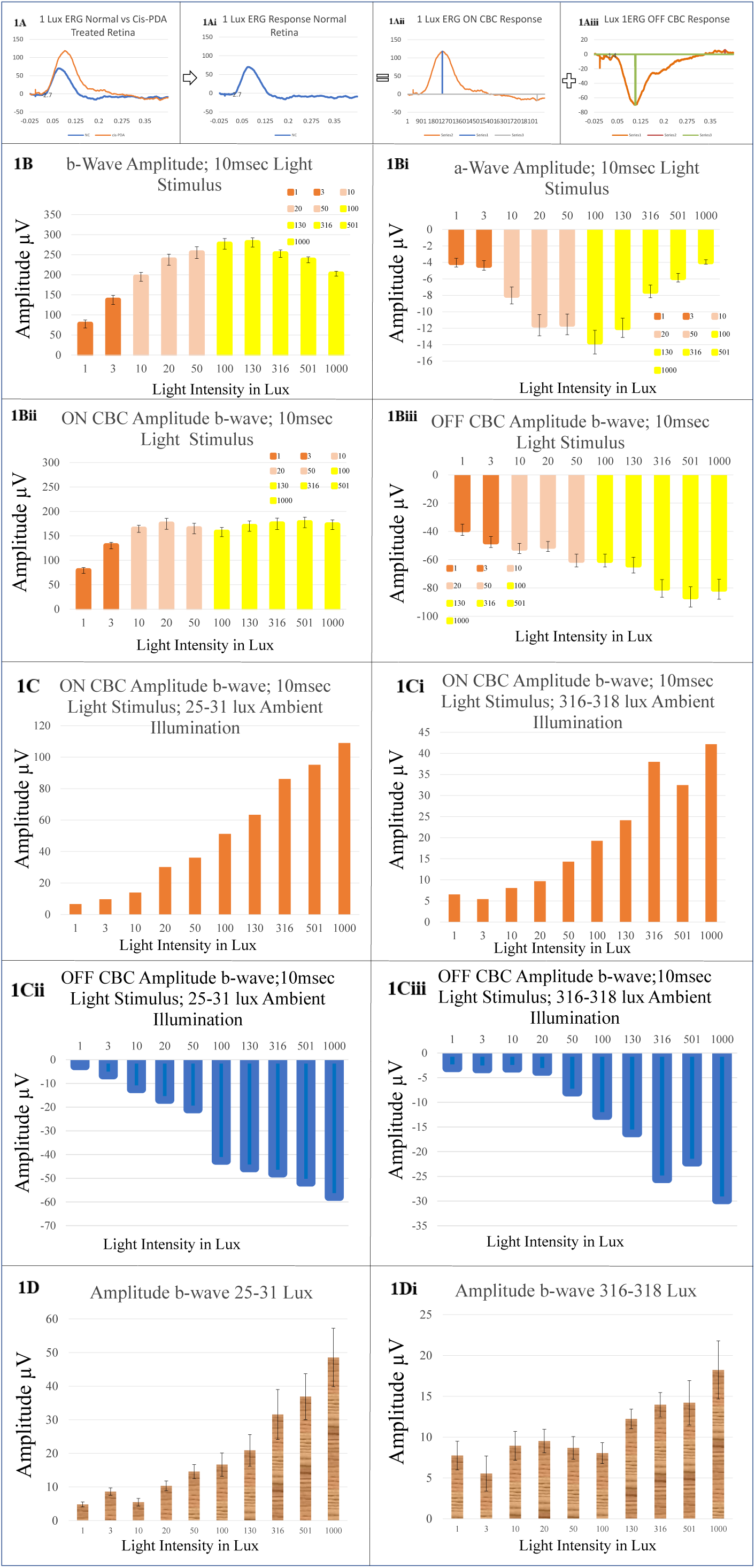
**A.** ERG recorded simultaneously against 1 lux white flash stimuli from both eyes of rat; one eye saline-treated control (response in blue) and other eye treated with 4µl 100mM cis-PDA (response in red). Subtracting cis-PDA response (1Aii: red) from normal ERG response (1Ai: blue) gives OFF cone bipolar response (1Aiii). **Fig. 1B**. ON and OFF cone bipolar response was isolated by subtracting cis-PDA response from the saline-treated eye against 1, 3, 10, 20, 50, 100, 130, 316, 501 and 1000 lux: one was eye treated with intravitreal injection of saline (pH 7.4) and other with cis-PDA. 1B: ERG b-wave and 1Bi: a-wave of amplitudes of control eye. 1Bii: ON cone and 1Biii: OFF cone bipolar amplitudes of cis-PDA treated eye. **Fig. 1C**. ON and OFF cone bipolar response of rat ERG using cis-PDA against 1, 3, 10, 20, 50, 100, 130, 316, 501 and 1000 lux at 25-31 lux (1C and 1Ci) and 316-318 lux (1Cii and 1Ciii) ambient illumination. **Fig. 1D**. b-wave response of normal eye ERG in ambient illumination: of 25-31 lux (1D) and 316-318 lux (1Di).

Subtracting this ON CBC response (Fig.1Aii) from normal ERG (Fig.1Ai) obtained from saline-treated contralateral (control) eye helped observe OFF CBC response to light stimulus which also increased with an increase in intensity of light stimuli (Fig 1Biii, Fig.1Cii and Fig.1Ciii) (Xu et al., 2003).

Xu et colleagues. investigated the influence of light intensity on both ON and OFF CBCs against a constant backdrop (to eliminate rod ERG) (Xu et al., 2003). However, there is no literature on the OFF CBC response to diverse ambient light conditions and light stimulation intensities. In the current work, we found ON and OFF CBC responses to increasing light stimulus strengths both in the absence (scotopic) and in the presence of lighted backdrops (mesopic and photopic). We discovered that the response of both ON and OFF CBCs increased with increasing light stimulus intensity in scotopic illumination in the dark-adapted retina (Fig. 1Bii & 1Biii). However, the response of the ON CBCs appears to saturate early with an increase in light intensity while OFF CBC response continued to increase consistently with greater light intensity (Fig. 1Bii & 1Biii).

### ON and OFF Bipolar response of rat ERG in ambient illumination

We also evaluated the effect of ambient illumination on ON and OFF CBC response of dark-adapted rats. In the initial study, we presented light flashes in the illuminating background of 25-31 lux (mesopic background, Fig.1C: ON CBC response & Fig. 1Cii. OFF CBC response). In a separate experiment, we measured ERG responses against an illuminating background of 316-318 lux (photopic background, Fig.1Ci: ON CBC response & Fig. 1Ciii. OFF CBC response).

Under scotopic illumination, ON and OFF CBC amplitudes increased up to ∼ 175 µV and ∼ 90 µV respectively and amplitudes of b-wave increased up to 290 µV (Fig. Bii, Fig. 1Biii & Fig. 1B.). At mesopic illumination (25-31 lux), ON and OFF CBC amplitudes increased to ∼115 µV and ∼ −60 µV respectively, and b-wave amplitudes increased to 50 µV. Under photopic illumination (316-318 lux), ON and OFF CBC amplitudes increased up to ∼40 µV and ∼ −30 µV respectively, and amplitudes of b-wave increased up to 19 µV (Fig.1C, Fig.1Ci, Fig.1Cii, Fig.1Ciii, Fig.1D and. Fig.1Di). Additionally, the b-wave response of saline-treated retina in dark-adapted conditions (Fig.1B) showed a typical pattern of increasing amplitude, saturation, and decline with stronger light flashes, while under bright light conditions, the response was generally exponential (Fig. 1D & 1Di.). In conclusion, both ON and OFF CBCs exhibited responses to increased light stimuli across all three ambient light conditions (scotopic, mesopic, and photopic), although the response magnitude diminished as background illumination increased, indicating light adaptation as summarized in Table 1.

**Table 1.** Comparison of ON and OFF cone bipolar response of rat ERG measured in scotopic, mesopic, and photopic illumination.

|  | ON CBC Response | OFF CBC Response |
| --- | --- | --- |
| Dark-Adapted | ↑↑↑ | ↑↑↑ |
| Mesopic | ↑↑ | ↑↑ |
| Photopic | ↑ | ↑ |

A plot of ON and OFF CBC amplitude response to light intensity also shows that for each lux increment, ON and OFF CBC amplitudes increase by 0.138 µV and 0.0427 µV respectively (Fig.2 Supplementary). Which means the response of ON CBC to each lux of light is approximately 3.23 times more than that of OFF Bipolar cells.

### Role of OFF CBCs to overall ERG waveform (OFF CBCs respond to Light mostly in an inhibitory manner) / OFF CBCs contribute to ERG mostly in an inhibitory manner

As mentioned earlier, we concurrently recorded ERG from saline-treated eye (normal ERG) and cis-PDA-treated (ON CBC response) contralateral eye of dark-adapted rat to different light intensities (ranging from 1-1000 lux) under scotopic conditions as per Xu et. al., 2003. We then extracted XY coordinates of normal ERG and ON CBC responses and deduced OFF CBC responses. Then, using the MS Excel program, we resynthesized the ERG waveform by adding XY values of ON CBC response at a certain lux level to the OFF CBC response at a certain other lux level using formulae ON CBC response + OFF CBC response = a resultant ERG. The purpose of doing this exercise was to understand what is the role and contribution of OFF CBC’s response to an ERG signal. We have summarised changes in ERG response observed if different strengths of ON and OFF CBC components are combined as given in Table 2 (also see Fig. 2A-2Fi).

**Table 2.** ERG response resynthesized combining ON and OFF CBC responses of different strengths obtained from dark-adapted retina under scotopic illumination. NR: Normal ERG (green); ↑ ERG: ERG increased (blue); ↓ ERG: ERG reduced (yellow); I ERG: Inverted ERG (red font)

| ON→<br>OFF↓ | 1 | 3 | 10 | 20 | 50 | 100 | 130 | 316 | 501 | 1000 |
| --- | --- | --- | --- | --- | --- | --- | --- | --- | --- | --- |
| 1 | NR | ↑ ERG | ↑ ERG | ↑ ERG | ↑ ERG | ↑ ERG | ↓ ERG | ↑ ERG | ↑ ERG | ↑ ERG |
| 3 | ↓ ERG | NR | ↑ ERG | NR | ↓ ERG | NR | ↓ ERG | ↑ ERG | ↑ ERG | ↑ ERG |
| 10 | ↓ ERG | NR | NR | NR | ↓ ERG | NR | ↓ ERG | ↑ ERG | ↑ ERG | ↑ ERG |
| 20 | ↓ ERG | NR | NR | NR | ↓ ERG | NR | ↓ ERG | ↑ ERG | ↑ ERG | ↑ ERG |
| 50 | ↓ ERG | ↓ ERG | ↓ ERG | NR | NR | NR | ↓ ERG | ↑ ERG | ↑ ERG | ↑ ERG |
| 100 | ↓ ERG | ↓ ERG | ↓ ERG | NR | ↓ ERG | NR | ↓ ERG | ↑ ERG | ↑ ERG | ↑ ERG |
| 130 | ↓ ERG | ↓ ERG | ↓ ERG | NR | ↓ ERG | NR | NR | ↑ ERG | ↑ ERG | ↑ ERG |
| 316 | I ERG | ↓ ERG | ↓ ERG | NR | ↓ ERG | NR | NR | NR | ↑ ERG | ↑ ERG |
| 501 | I ERG | ↓ ERG | ↓ ERG | ↓ ERG | ↓ ERG | ↓ ERG | NR | NR | NR | ↑ ERG |
| 1000 | I ERG | ↓ ERG | ↓ ERG | ↓ ERG | ↓ ERG | ↓ ERG | NR | NR | NR | NR |

**Fig. 2.**
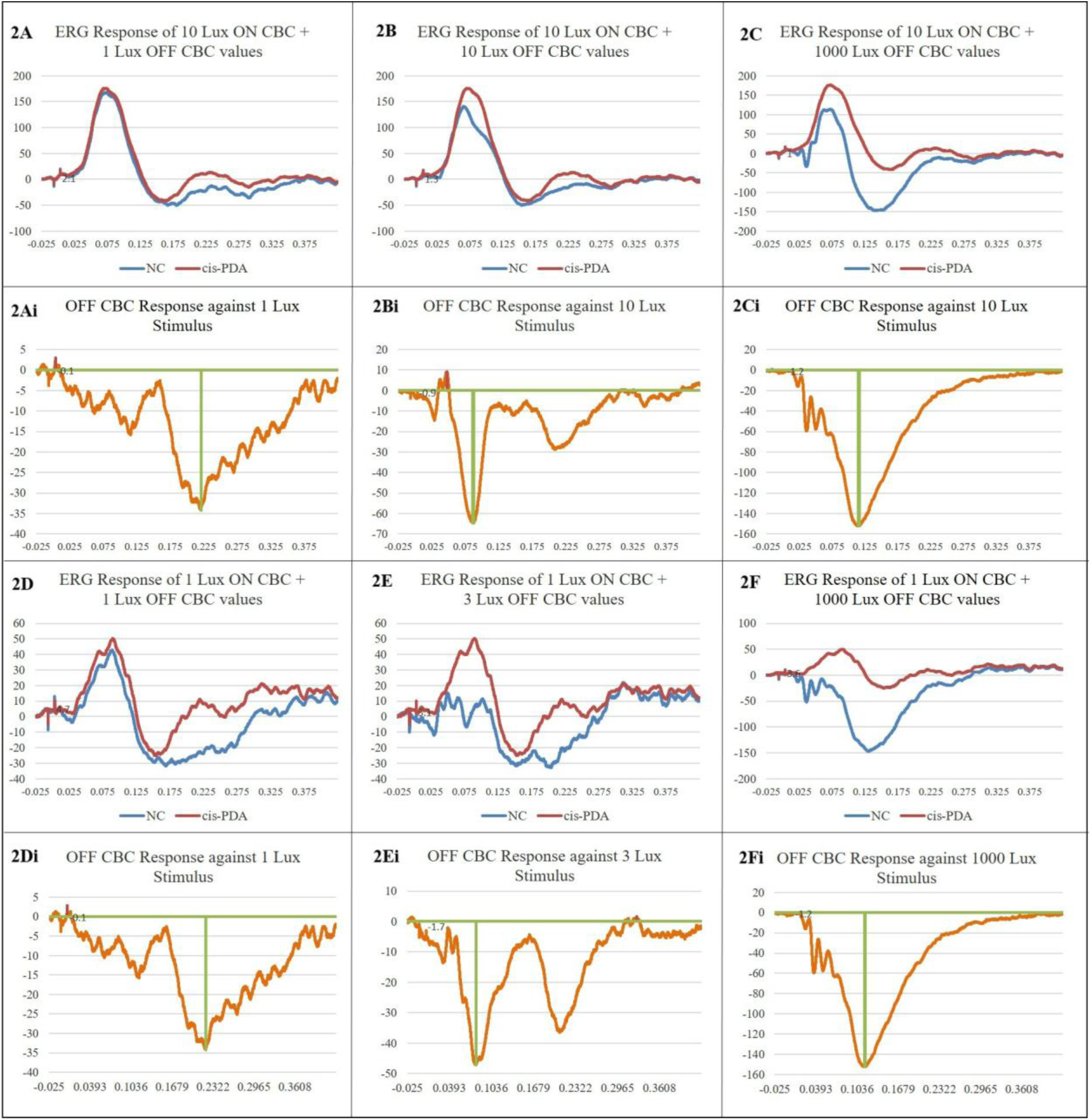
**A.** ERG response to 10 lux stimuli (2B) vs change in ERG response when OFF CBC response is reduced (equivalent to 1 lux stimulus) (2A) vs change in ERG response when OFF CBC component is increased (equivalent to 1000 lux stimulus) (2C). Note when the OFF CBC is significantly reduced, the overall ERG (blue) response increases (2A & 2Ai), and when the OFF CBC is significantly increased overall ERG response (blue) is significantly reduced (2C & 2Ci). Fig. 2D. ERG response to 1 lux stimuli vs change in ERG response when OFF CBC response is increased (equivalent to 3 lux stimulus) (2E) vs change in ERG response when OFF CBC response is increased (equivalent to 1000 lux stimulus) (2F). Note as the OFF CBC response increases overall ERG (blue) response reduces and even becomes inverted when the OFF CBC response increases significantly (2D-2F). Blue trace: Overall ERG response; Red trace: Response of ON CBC and orange trace: OFF CBC response.

Our findings reveal that when contribution of OFF CBCs exceeded that of ON CBCs, the overall ERG was either diminished or inverted. This phenomenon was particularly evident at low light intensity stimuli (1-10 lux). In contrast, when the contribution of OFF CBCs was lower than that of ON CBCs, the overall ERG increased (resulting in a supranormal ERG) compared to the anticipated normal ERG response (marked in green). This observation was most prominent at higher light intensities (316-1000 lux). This suggests that variations in OFF CBC contributions influence the shape of ON CBC responses, thereby affecting the overall ERG signal. At light stimuli of 20 and 100 lux (mesopic range), OFF CBCs appeared to contribute minimally to the overall resultant ERG response, yet the pattern was consistent; meagre OFF CBC contribution led to an elevated ERG while a very high OFF CBC contribution resulted in diminished ERG. Therefore, it can be concluded that OFF CBCs modulate ON CBC response to light through some inhibitory mechanism. However, at 50 and 130 lux, the contribution of OFF CBCs to overall ERG was a bit unusual and even a small drop in OFF CBC response caused a reduction in resultant ERG instead of an increase. We observed mixed results at this light intensity (also pertaining to a mesopic stimulus) that will need further investigation (Table 2).

To understand the significance of OFF CBC’s response to light, we re-analyzed retinal architecture (Fig. 3A) and its orientation to eye orbit. Anatomical and microscopic studies suggest retinal bipolar axons project parallel to the plane of light and their terminals innervate the inner plexiform layer (IPL) at two different levels: Sublamina a and Sublamina b. OFF CBC terminals ramify in Sublamina a (making 40% IPL thickness) and ON CBC terminals ramify in Sublamina b (making 60% IPL thickness) (Strettoi et. al., 2010) and terminal arbors of both cell types branch perpendicular to the plane of light popularly known as “Topological rule of retinal architecture” (Famiglietti et.al., 1997; Zhang et. al., 2021).

**Fig. 3.**
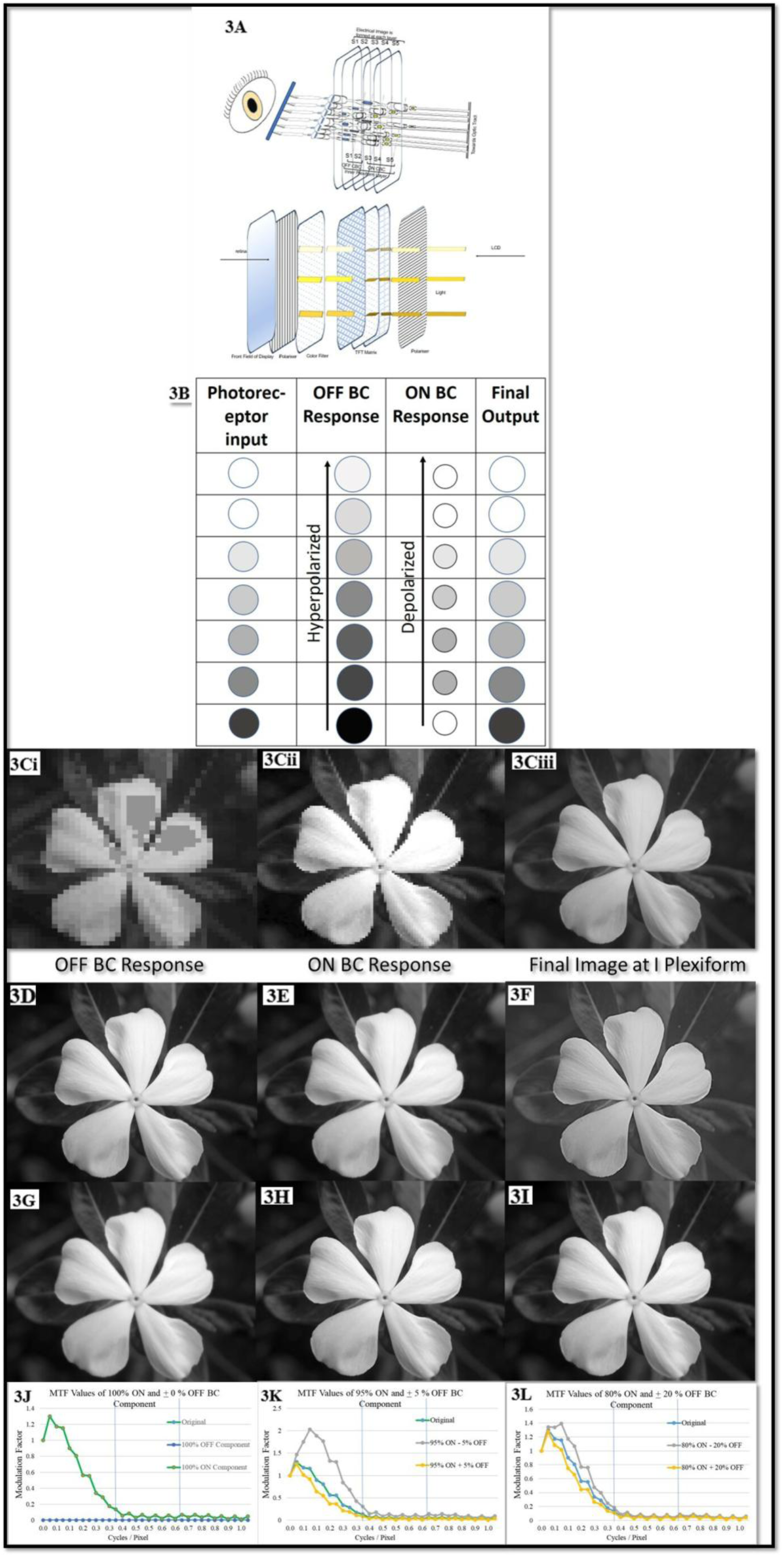
**A.** Representative image of the orientation of retinal neurons inside eye orbit. Two screens of OFF CBCs lie behind 3 screens of ON CBCs. The electrochemical signal from the photoreceptor terminal (circle with arrow) is simultaneously transmitted through both ON and OFF CBCs. Both OFF CBC and ON CBC electrical image components of varying voltage gradients are formed at 5 sublamina’s. **Fig. 3B**. Our representation of how photoreceptor output is bifurcated into ON and OFF CBC electric field images which amalgamate into a final output to the brain. **Fig. 3C**: Representation of an ON CBC response (3Cii) superimposed on an OFF CBC response (3Ci) to give a clear and sharp image of a vinca flower (3Ciii). **Fig. 3D**. Combination of ON and OFF CBC images: 3D: Original image (100% ON + 0% OFF), 3E: 95% ON + 5% OFF CBC, 3F: 95% ON - 5% OFF CBC, 3G: 40% ON + 60% OFF CBC, 3H: 110% ON + (−10) % OFF CBC, 3I: 110% ON – (−10) % OFF CBC. **Fig. 3J**. MTF graphs of output images obtained by adding ON and OFF BC image where contribution of ON bipolar cells is more than that of OFF bipolar cells: Original, 100% ON and 0% OFF BC; 3K: Original, 95% ON ± 5% OFF BC, 3L: Original, 80% ON ± 20% OFF BC.

This means that dendrites of these CBC layers ramify such that OFF CBC terminals form some sort of electrical screen/web posterior and parallel to the screen formed by ON CBC terminals. In other words, the OFF CBC electrical image screen lies closer to photoreceptors and the ON CBC electrical image screen lies closer to retinal ganglion cells. All the screens (S1-5) lie parallel to each other. Further, electrical signals from photoreceptor cells converge at I plexiform such that visual information is transmitted pixel-wise into these 5 parallel sublaminae’s. Therefore, I plexiform appears as optical equivalent of a TFT screen of an LCD. As with photoreceptor cells, both foveal density as well as eccentricities of these CBCs increase towards the fovea suggesting extensive overlap (Zhang et. al., 2021) leading to the formation of a finer electrical image at the central retinal region compared to the periphery.

We, therefore, conducted synthetic image experiments to investigate the role of OFF CBCs in creating light-associated electrical images at I plexiform, considering the fact that OFF CBCs’ response strength is less than that of ON CBCs’ response to the same light intensity stimuli and their density is also lower. In other words, we aimed to understand how an object, image, or dot of light would be perceived at OFF CBC level, which have a 30% density at I plexiform, compared ON CBC level which have 70% density.

### Nature of Image Modification by OFF CBCs: MTF Analysis

We, therefore, used a clear monochrome image of the vinca flower and labeled it as an “ON image” (by ON CBCs). Then we used Image J software to use monochrome type 8bit Gaussian Blur of the same image as “OFF image” (by OFF CBCs). We then imported numeric pixel-wise image values of both ON and OFF CBC components, and mathematically added/subtracted different percentages of OFF CBC image values from different percentages of ON CBC image values (Table 1. Supplementary). The resultant numeric pixel-wise output image file was then imported back into Image J and converted into ‘png’ format (Fig.3D-3I.).

A Mathematical transfer function (MTF) analysis of all combinations of ON and OFF CBC images was also carried out (Fig. 3J-3L) to check whether the OFF CBC component adds to or subtracts from the ON CBC component to give a better or sharper output image. And what is the likely mathematical contribution of the OFF CBC component to the ON CBC component that gives a sharper and clearer image. We found a 5% blurred-off CBC pixel-wise numerical image when subtracted from a 95% pixel-wise numerical ON CBC image gave the sharpest output image where edges could be detected most clearly. This means OFF CBCs shape response of ON CBCs to light intensity majorly by some inhibitory effect and this helps in sharpening of an electrical image at I plexiform.

## Discussion

### Response of Photoreceptor Cells to Light

It is well-known that in the retinal circuitry, photoreceptor cells function as photon detectors, responding by membrane hyperpolarization which reflects as “a-wave” in an ERG signal (Fig. 1Bi). The mechanism by which photoreceptor cells react to subtle variations in light intensity and wavelength has been extensively studied (Pugh and Lamb 2000) and are consistent across species. These cells can be regarded as photodiode transducers that modulate neurotransmitter (glutamate) release in response to light. At I plexiform, photoreceptor signals are known to bifurcate into ON (through rod bipolar cells and ON CBCs) and OFF (through OFF CBCs) pathways (Fig.1B-1Biii and Fig.1C-1Ciii). Retinal photoreceptors are known to release glutamate in the dark and this release diminishes with increase in light intensity. A rise in glutamate levels during the dark depolarizes OFF CBCs and triggers OFF-centre ganglion cells (OFF GCs) to fire. Conversely, glutamate reduction during illumination results in ON CBC depolarization which triggers ON-centre ganglion cells (ON GCs) to fire.

Therefore, it is interpreted that a central illumination and surround inhibition cause ON CBCs to depolarize leading to increased firing of ON GCs. While surround illumination and central inhibition cause depolarization of OFF CBCs leading to increased firing of OFF CBCs. This difference in responses between ON CBCs and OFF CBCs under varying illumination conditions result in contrast detection, which is a fundamental aspect of visual perception of differentiation in light intensities or colours (wavelength’s) between two adjacent spots. This mechanism plays an important role at the cellular level and influences how we perceive our surroundings visually in terms of textures or patterns.

However, this does not explain why OFF GCs are not firing when there is no illumination in both the center and surround because OFF CBCs at the center would be expected to depolarize. Also, why do OFF GCs fire when the center and surround are illuminated as OFF CBCs are not depolarized? Because by definition these cells should ideally be inhibited by light at the centre. Further, why do OFF GCs respond with low frequency firing equivalent to ON GCs firing due to excitation by light in the surround? (Fig. 4, Table 3.).

**Fig. 4.**
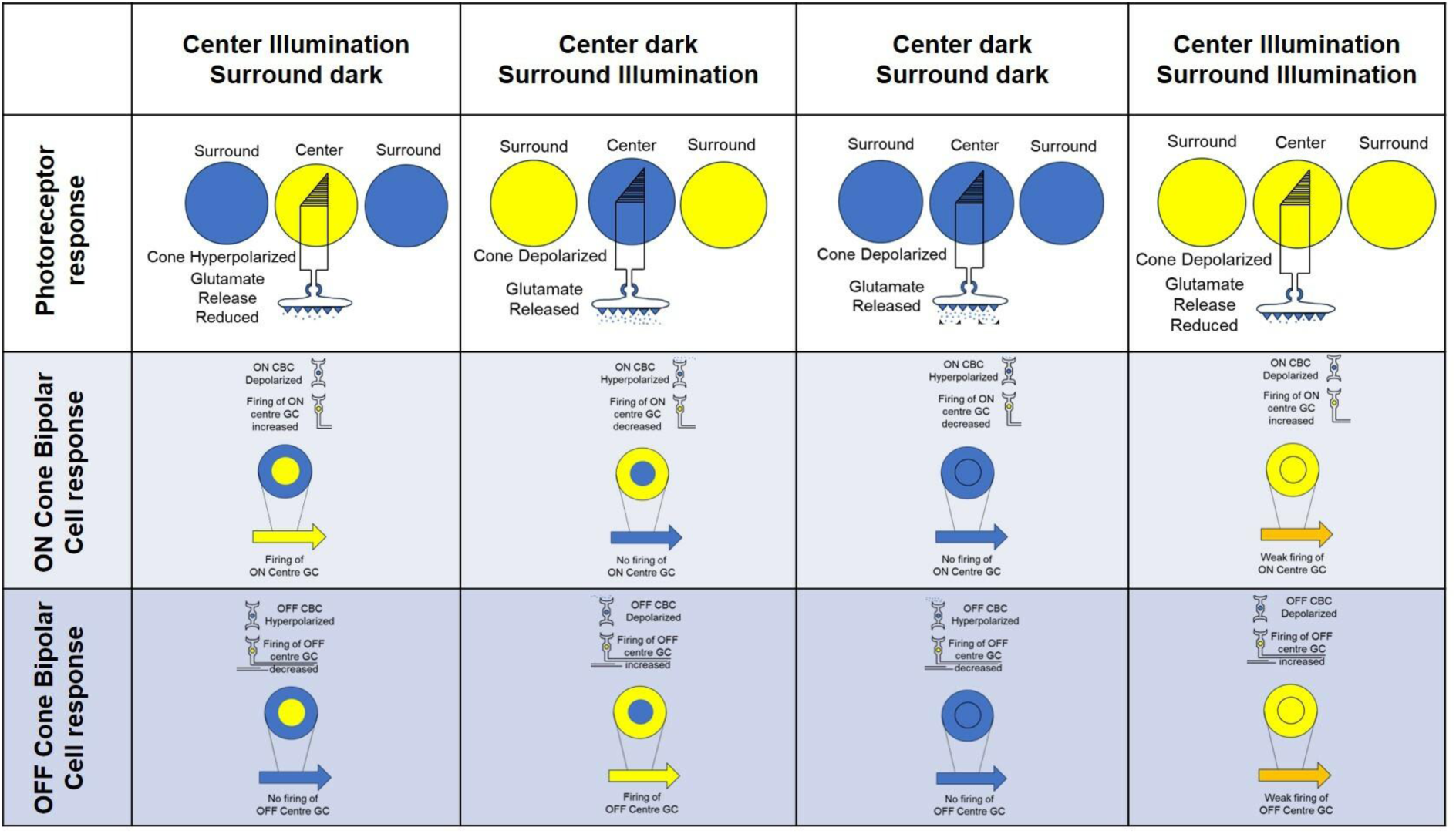
Concept of Luminance detection in the retina: Light photon triggers an electrochemical signal at the photoreceptor-bipolar synapse. Which causes increased firing of ON CBC and associated ON GCs. Light illumination (shown as yellow circle and arrow) at the centre or surround causes more firing of associated ON GCs in the field (blue circle and arrow represents dark). The difference in the firing rate of two adjacent ganglion cells helps determine the contrast between the two points.

**Table 3.**
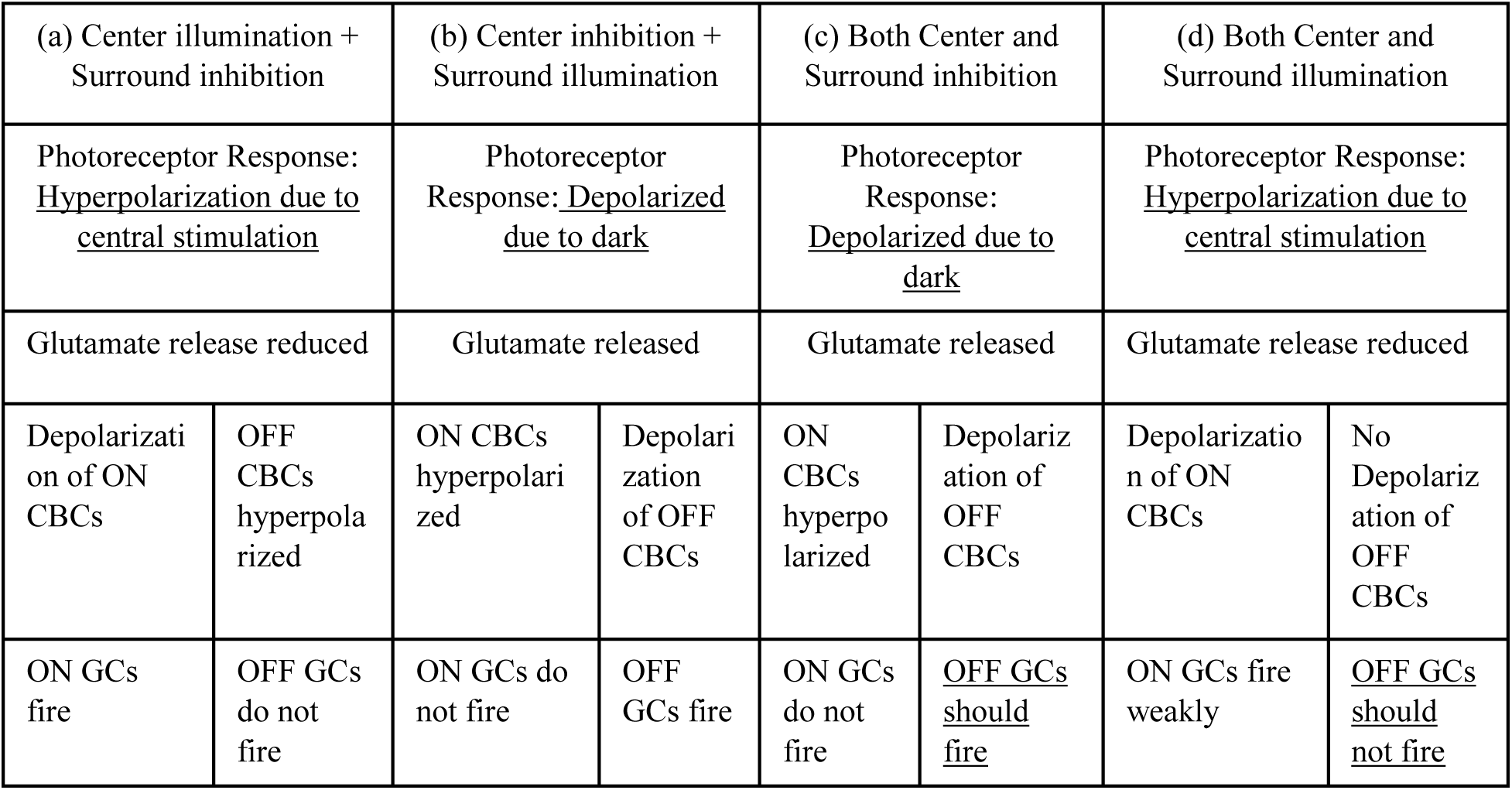
Response of ON and OFF CBCs to (a) Center illumination + Surround inhibition, (b) Center inhibition + Surround illumination, (c) Both Center and Surround inhibition and (d) Both Center and Surround illumination.

We therefore aimed to probe the role of retinal bipolar cells in visual image transmission. In the light of published reports and results of the present work, our interpretation is as follows:

### OFF Bipolar Cells also Respond to Light Signals and contribute to image formation at I plexiform

We hypothesized that OFF CBCs also contribute to electrical image transmission at I plexiform and therefore an important component of b-wave. So, we measured responses of both OFF and ON CBCs to increasing light stimulus intensities (Xu et. al., 2003) and under ambient light conditions (Mojumder 2008) (Fig. 1A-1D series). We found:

(1) **Response of both ON and OFF bipolar cells increased with light stimulus intensity**, suggesting bipolar cells of both rod, cone, as well as mixed rod + cone pathway (refer Table 4 / Fig. 1A), respond to light-induced glutamate reduction at I plexiform. Our results are consistent with earlier reports (Xu et. al., 2003). In the present study, we also found that OFF CBCs give response to the light stimulus presented in scotopic illumination, which agrees with earlier reports from cellular studies (Hack et al., 1999; Li and DeVries, 2004; Strettoi et. al., 2010) that OFF CBCs receive signals from rods. Therefore, we conclude that light-associated electrical signals pass via the rod bipolar + OFF CBC pathway in dim light (scotopic illumination).

**Table 4.**
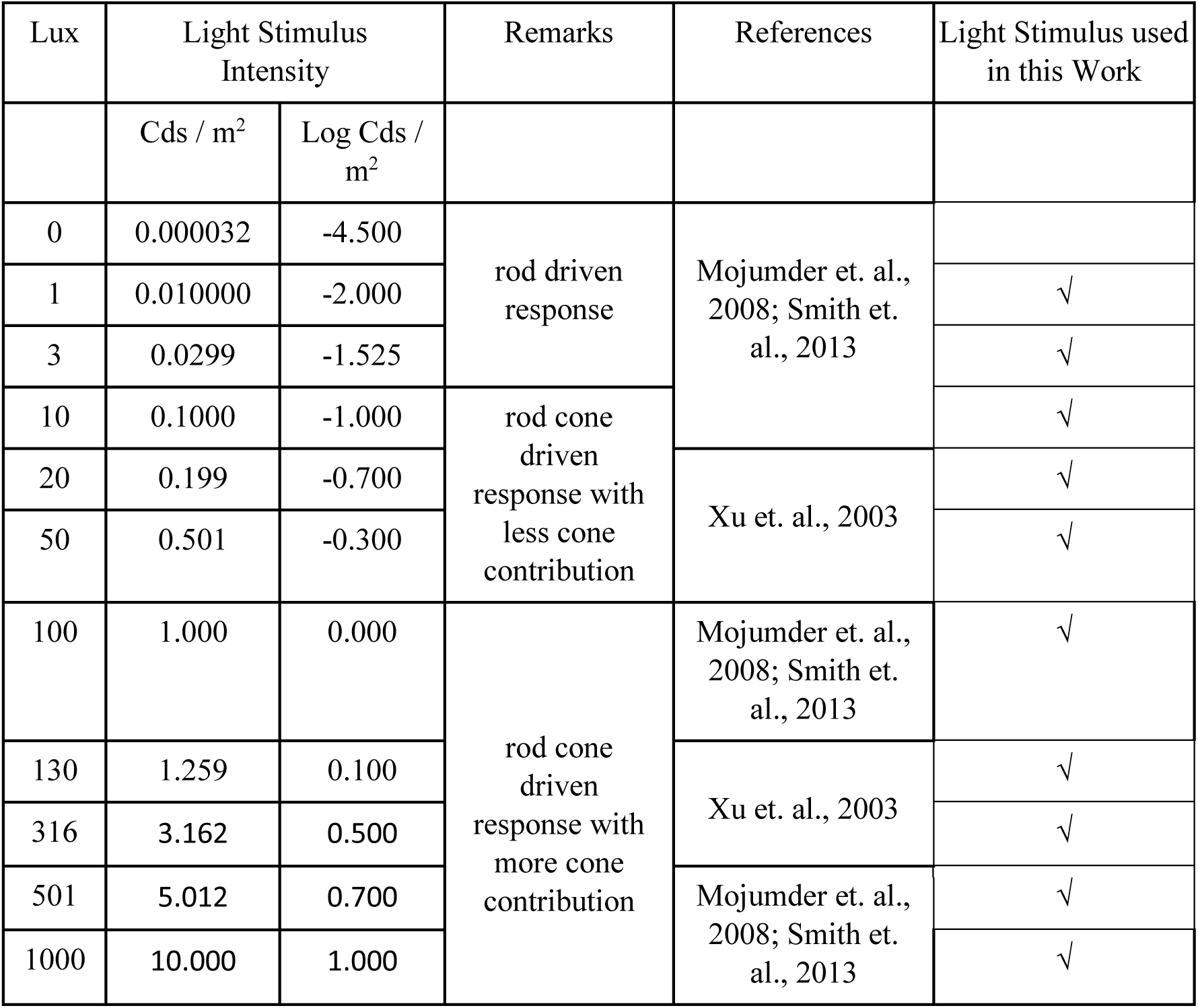
Different stimulus strengths used in the present study to excite rod driven, mixed rod cone driven response with less cone contribution.
(2) **Response of both ON and OFF CBCs increase with intensity of light stimuli in both dim as well as ambient light conditions, although their overall amplitudes reduce with increase in intensity of background:** We found that for same pre-synaptic stimuli (light stimulus of 1-1000 lux), ON CBCs depolarized up to 138 counts (82-220 µV; higher range) while OFF bipolar cells hyperpolarized up to only 43 counts (from (−42)-(−85) µV; lower range) under scotopic conditions. Under mesopic illumination (25-31 lux), these values were 6.5-108 µV; 101.5 count range and (−3) to (−58) µV; 55 count range for ON and OFF CBCs respectively. These values reduced further to 6.5-42 µV; 35.5 count range and (−3) to (−30) µV; 27 count range for ON and OFF CBCs in photopic illumination (316-318 lux) (Fig. 1C-1Ciii & Fig. 1D-1Di). Suggesting both ON as well as OFF CBC respond linearly to light stimuli irrespective of ambient illumination. With increasing ambient lighting, the ratio of ON bipolar to OFF CBC response (3.209 in scotopic illumination vs. 1.854 in mesopic illumination vs. 1.5 in photopic illumination) continues to decrease, indicating an increased or similar contribution of OFF CBCs under settings that are suited to light. Our results align with cellular studies that show glutamate reduction (at I synapse) causes simultaneous depolarization of ON CBCs and hyperpolarization of OFF CBCs. Wherein, ON CBCs depolarize to a degree of 20 to 55 counts (Awatramani and Slaughter, 2000; Euler and Masland, 2000), whereas OFF bipolar cells hyperpolarize to a degree of 7–16 counts (Euler and Masland, 2000), despite having similar resting membrane potentials. **This indicates both ON and OFF CBCs operate similarly diodes converting neurotransmitter signal at I plexiform into graded electrical responses at II plexiform and this response property of CBCs appears to be universal. It is worth noting that the response of OFF CBCs correlates directly with an increase in stimulus input and this response attribute is comparable to photoreceptor or ON CBCs** (Fig. 1Bi, Fig. 1Biii, Fig. 1Cii & Fig. 1Ciii) **indicating that these cells are also involved in creation of electrical images at I plexiform**.

### OFF Bipolar Cells fine-tune and thus modify electrical image formed by ON Bipolar cells at I plexiform

The present study revealed that ON CBCs subtly responded to increasing light intensity under scotopic illumination that saturated earlier than a typical b-wave response (Fig. 1Bii). But OFF CBCs displayed exponential response, saturating considerably later (Fig. 1Biii). Nevertheless, the overall b-wave manifested as a progressive depolarization until reaching a saturation, followed by a decline (Fig. 1B). It is proposed that OFF CBCs alter how ON bipolar cells respond to light in scotopic, mesopic, and photopic illumination (Figs. 1C, Fig. 1Ci, Fig. 1Cii and Fig 1Ciii). It is also proposed that a ratio between ON and OFF CBC response is essential for normal ERG, which when disturbed results in either a supranormal, reduced, or even inverted ERG (Fig.2A-2F and Table 2). Such atypical ERGs have been clinically noted in numerous retinal disorders associated with abnormalities (such as photophobia) or even vision loss.

Our image analysis (MTF; using Image J software) studies corroborate OFF CBCs generate a minor but significant dark and blurred image which when subtracted from the ON image, produces a final clear image that surpasses the clarity achieved at the level of ON CBCs alone. Thus, a modest contribution from OFF CBCs helps refine the overall electrical image at I plexiform through unsharp masking enhancing image sharpness via edge detection thereby contributing to visual acuity.

### Visual image at I plexiform is formed in electric field layers of OFF and ON cone bipolar cells

Our observations and conclusions align with our view of retinal architecture (Fig. 3A) and its orientation to eye orbit. Wherein retinal bipolar axons terminals project parallel to the plane of light to form electrical image screens at five different sublaminae’s (Strettoi et. al., 2010, Zhang et. al., 2021). It is proposed that electrical image is formed layer-by-layer in inner plexiform which converge and combine to yield a composite image at level of ganglion cells.

The structure and operation of this retinal architecture resemble that of an LCD. Here an image sensor (composed of a grid of photo sensors) produces a charge when struck by light (akin to photoreceptor cells). Generated charges are then transformed into a series of continuous analog voltages at the output terminals, akin to retinal bipolar cell terminals. These terminals extend toward their branched dendritic ends at right angles. At each level of the sublamina, these electric fields combine pixel by pixel to form an amalgamated electrical image. An electric field continuum is established by the fusion of five such amalgamated electrical images that arise at each sublamina: two layers representing OFF CBCs (low amplitude hyperpolarizing occupying 40% IPL thickness) and three levels representing ON CBCs (high amplitude depolarizing occupying 60% IPL thickness). This means that by blending various proportions of OFF and ON CBC outputs, the outer plexiform layer (OPL) can process light of different intensities (Fig.3B).

### Blurred Visual image in OFF electric field layers contributes to unsharp masking

Our synthetic image experiments using Image J analysis reinforce our conclusion that a blurred low-intensity OFF CBC image (due to low density and amplitude) combines with a bright high-intensity ON CBC image (due to high density and amplitude) to yield an overall sharp vision. When the OFF CBC electrical image is subtracted from the ON CBC electrical image, the resultant image is clearer than the one produced at ON CBCs level (Fig.3Ci-3Ciii). Such mechanism could enhance sharpness by rectifying high intensity fluctuations especially along edges of the image, thus improving vision. This enhancement is possible because the OFF CBCs form a small yet considerable dark and blurred image.

### Significance of our work

Despite vision research starting at least a century earlier than auditory research, electrical devices to restore hearing were invented in 1900s while an analogous electrical stimulation of retina to restore vision is yet to be achieved. This challenge is due to our insufficient understanding of signal processing within retinal network and necessitates re-analysis of existing literature on retina.

Current literature suggests photoreceptor cells stimulate downstream CBCs when struck by photons. CBCs in turn transmit signals to the corresponding GCs. This means that retinal ganglion cells fire when light enters their receptive field (Fig. 4). A central illumination means an increased firing of ON GCs and concurrent decreased firing of OFF GCs at the centre. And vice versa with surround illumination. This difference in firing helps in contrast detection. However, this interpretation does not help understand weak firing response (due to mutual inhibition) of both ON and OFF GCs when both centre and surround are illuminated when ideally, electrical fields should summate/interact with each other through the principle of superposition.

We propose that an increased firing of ON or OFF GC is directly proportional to the signals received from corresponding ON and OFF CBC sets in response to a particular light spot. That is, a light spot of high intensity will generate more response from ON CBCs, and cause more firing of ON GCs. And a light spot is of low intensity will cause more OFF GCs to fire. It is worth noting that GCs are also arranged as layers akin to CBCs. In certain mesopic intensities, both GCs would fire similarly. Therefore, if the two adjacent light spots are of the same intensity, both types of GCs would fire unvaryingly. Our interpretation helps explain and understand “weaker responses” when both the centre and surround are illuminated. We envisage so because when two bright screens lie alongside, the common edges between them would be brighter. But when they lie behind another, both their brightness is somewhat negated, edges sharpened and contrast created (Fig. 5). This part will however need to be experimentally verified.

**Fig. 5.**
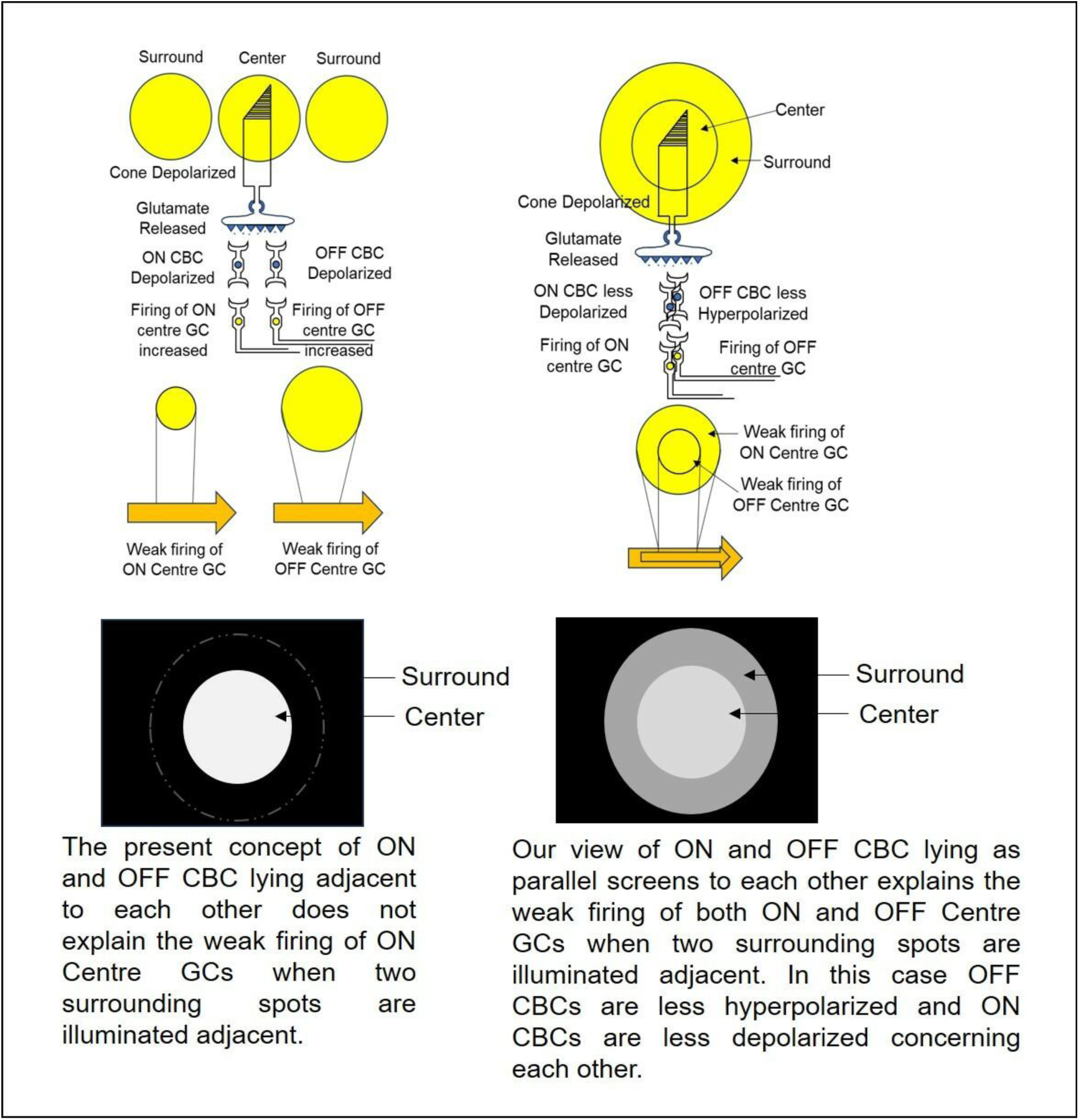
Our interpretation of electrical image formed at I plexiform. Left Top: Image formed when ON and OFF CBC responses are alongside each other. Left Bottom: How images formed due to two light spots when ON and OFF CBC responses are alongside each other as per current prevalent opinion. Right Top: Our representation of an image formed when OFF and ON CBC responses are oriented as parallel screens. Right Bottom: when these responses is seen in space.

We therefore propose that electrical signals at photoreceptor terminals diverge into parallel ON and OFF CBC screens at I plexiform, wherein each CBCs dendrites ramify perpendicularly, producing a little voltage spread equal to the light “spots” intensity. Signal summation is the outcome of these distinct voltage distributions most likely overlapping at common locations where dendritic arbours connect. Consequently, this mechanism helps the retina process a wide range of light intensities across the visual field. A sharp 2-dimensional vision is also produced by electrical signals that summate parallel across sublamina. Because OFF CBCs help to create this crisp electrical image by shaping light intensity response at IPL. Undoubtedly, 40% of IPL thickness occupied by OFF CBCs which contribute to 30% of ERG b-wave will not be without any purpose.

Similar conditions are found in nature. Raindrops are translucent; thus, we cannot see them until they have a boundary and a background! Like translucent glass, the medium in which we live—mother nature, the environment, and space—has no background. Thus, we may see through it to observe the things (trees, animals, buildings, etc.) that have both an edge and a background. Background signals may be exemplified by other physiological systems. The QRS complex in an electrocardiogram (ECG) represents depolarization of neurons supplying ventricles that causes its contraction signifying blood flow throughout the body. However, a negative polarity Ta wave that represents atrial repolarization occurs simultaneously at a point matching to the QRS complex which is crucial because it indicates atrial blood refill for the successive cardiac cycle, even though it is obscured by the QRS complex. Without a Ta wave, there would be no atrial filling and, consequently, no blood being pumped into the systemic circulation from the ventricles. Thus, a concealed signal is a significant background activity to a big signal, in this example a QRS complex.

**We therefore propose OFF CBCs serve as nature’s canvas (coarse background) over which fine image details from ON CBC are etched out. An image of OFF CBC alone is quite dull and blurry while an image of ON CBC might come as dazzling streaks. But when both images are clubbed (the OFF component is mathematically subtracted from the ON component), the final output is a clear precise image.**

#### Potential Physiological relevance of current work

It is worth noting that despite similar RMP (−40mV and −36mV), light stimulus causes simultaneous graded responses from both ON and OFF CBCs albeit of opposite polarity and different magnitudes. This response property is universal amongst species although there could be some variation in their response sensitivities (similar to skin, hair color or heart rate, blood pressure, etc). Therefore, few people are more sensitive to bright light than others such that they cannot sometimes open their eyes. Similarly, an object might look brighter for a few people.

What if this range of normal response is altered? We look at a few possibilities as follows (Fig.6):

1. Case A: Less ON CBC response to light compared to OFF CBC: Hazy / Blurry and dim vision with reduced ERG b-wave response;
2. Case B: Negligible ON CBC response to light compared to OFF CBC: Loss of vision in dark with Inverted ERG;
3. Case C: No ON CBC response to light / poor OFF CBC response: A sudden decrease in vision even during daytime, seeing what looks like a curtain coming down over one eye with very reduced / negligible ERG b-wave response;
4. Case D: Occasional ON CBC response / No OFF CBC response: Seeing flashes of light, seeing halos or rainbows around lights (like a lightning) with occasional b-wave response could result in photophobia;
5. Case E: Hypersensitive response of ON and OFF CBC to light: Sudden sensitivity to light and glare, seeing floaters or spider webs could also result in photophobia;
6. Case F: Loss of ON or OFF CBC at peripheral retina: Tunnel vision. Two types of tunnel vision can be there: central part is clear (ON and OFF CBC working) + (a) Peripheral part has no ON and OFF CBC response: vision limited to center and nothing at periphery OR (b) Peripheral part has OFF CBC response only: vision limited to center and it is like curtain coming down at periphery;
7. Case G: If impulses to OFF and ON CBC do not arrive at the same time/latency of ON CBC is more than OFF CBC: Double vision (something like arrhythmia of heart or epilepsy of neurons; irregular or sudden discharges); 8. Case H: No response of either ON or OFF CBC: Blindness. We deduce this because people who are born blind neither see light nor dark. They see absolutely nothing.

**Fig. 6.**
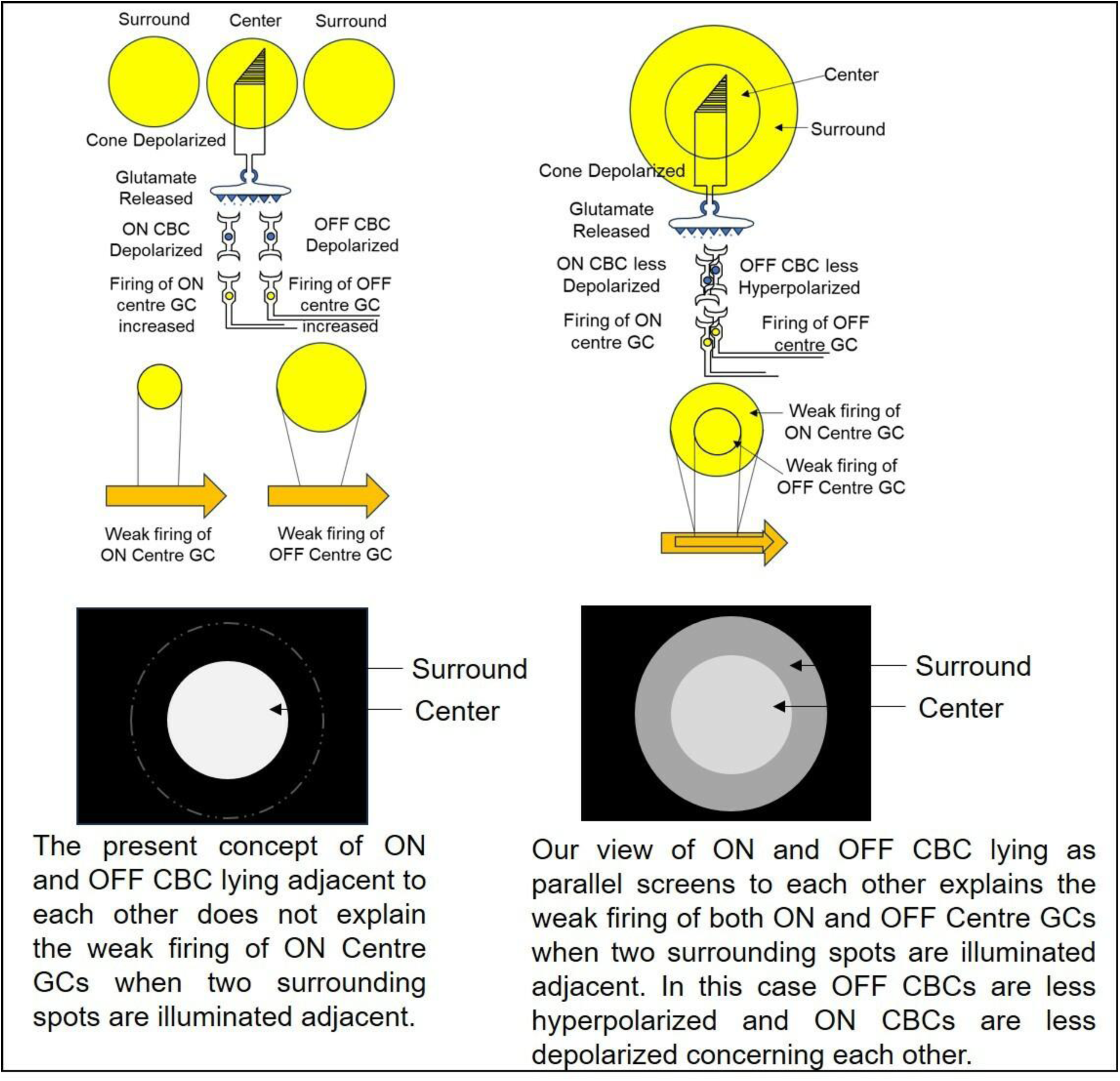
Different vision abnormalities explained with our model. A: Image formed when ON CBC (white) superimposes over OFF CBC image (this image will be used as reference to imagine component of OFF and ON CBCs. B: Less ON CBC response to light compared to OFF CBC; C: Negligible ON CBC response to light compared to OFF CBC (inverted ERG); D: No ON CBC response to light / poor OFF CBC response; E & F: Occasional ON CBC response / No OFF CBC response or Hypersensitive response of ON and OFF CBC to light; G: Loss of ON or OFF CBC at peripheral retina (Tunnel vision) with peripheral retina having no ON and OFF CBC response or peripheral retina has OFF CBC response only; H: Double vision when OFF and ON CBC images do not arrive at the same time; I: Complete blindness with no ON and OFF CBC response.

#### Potential Clinical relevance of current work

In the current study, we discovered that the b-wave in ERG may be lowered in one of two ways: when the ON CBC response component is reduced or when the OFF CBC response is proportionally greater than the ON CBC response. We also discovered (by image analysis) how electrical images created at I plexiform can be altered when different strengths of ON and OFF CBC images are combined. This is the first time a model has been presented that describes the nature, function, and cause of OFF CBC cells. Using this paradigm, we can explain not just the physiological contributions of OFF CBCs, but also the pathophysiology of visual impairments reported in many retinal disorders. We feel that our logic helps us understand the many visual abnormalities caused by improper signaling at I plexiform. Thus, we argue that differential measurement of ON and OFF pathway activity in humans will aid in more precise disease detection and treatment monitoring (Norcia AM et. al., 2020).

#### Potential Biomedical application/relevance of current work

The present study is the first to investigate the role of OFF CBCs in image formation within the retina. The current study suggests that because OFF CBCs contribute to ERG b-wave and therefore to electrical image formation, it will be worthwhile to stimulate these cell types in an appropriate ratio to restore even a modicum of vision. If we have a technique of transducing light energy (photons) into electric charges that causes a change in voltage of a semiconductor material. And if we can construct a linear relationship in such a system so that an increase in light intensity results in a proportional increase in semiconductor voltage, we will be able to develop an electrical artificial component of the retina’s IPL. This can be utilized to stimulate retinal ganglion cells converting analog I plexiform impulses into digital II plexiform signals. Which can then be forward propagated to the visual cortex via the optic nerve. Such systems exist. The photoelectric effect has been utilized in electronic devices designed for light detection. If such a photoelectric material is combined with material that shows photovoltaic capabilities, it will be possible to reproduce I plexiform functioning.

#### Future Scope of Work

Many different sizes of retinal bipolar cells have been identified. However, their exact innervation to various sublamellas of I plexiform, as well as their size, diameter, and response properties (in voltage) are not well understood. A good retinal bipolar isolation technique coupled with antibodies to the cell surface would give more precise and detailed information on the size and dendritic spread of each bipolar cell under a suitable microscope. When this information is combined to histological information from the retina, an equivalent semiconductor-based circuit can be designed to convert a photographic image into an electrical image. We propose information on ratio of bipolar cells employed to process each pixel of retinal image can be employed to make a sub-retinal artificial retina implant chip that can be utilized to restore vision in blindness due photoreceptor degeneration.

## Conclusion

Our results suggest both ON and OFF CBC contribute to two-dimensional vision, as light intensity results in a graded response to light by both types of bipolar cells. This information will be important in understanding the pathophysiology of various retinal diseases through differential assessment of ON and OFF CBC responses to light. Furthermore, tactics that proportionally activate both bipolar cell types will be beneficial in restoring vision since it will help the formation of sharp clear electrical images in the retina. Such an approach can be applied to develop electronic chips that operate as both ON and OFF Bipolar cells, effectively replacing outer plexiform and stimulating retinal ganglion cells to restore some functional vision.

There are many entities in this world that cannot be seen by human eye since human eye can detect only visible spectra. But it is well known that animals (butterflies, rats, dogs and birds) can see in ultraviolet or infrared spectra. If we can capture images in this spectrum and convert them into equivalent light stimulus that can trigger ON and OFF CBCs, we should be able to perceive a flower in real time (for example) from eyes of a mosquito, or a bee or a bird using the same prostheses, implants, or special glasses.

We believe our analysis of CBC response provides a novel perspective and a much more logical explanation of physiology of visual signal processing at I plexiform. And provides a better insight to contribution of RBCs to two-dimensional image formation at I plexiform level. We hope that our improved understanding of the roles of both these bipolar cell types in photon signal processing may aid medical practitioners in better understanding of retinal diseases that impair image formation. And will also help researchers and engineers worldwide to develop a new strategy and means of stimulating neural network at I plexiform through sub-retinal implants in order to restore vision.

## Acknowledgments

This work was supported by a BioCARe grant from the Department of Biotechnology (No. BT/PR19955/BIC/101/589/2016), Ministry of Science & Technology, Government of India. We thank Dr. Yogesh A. Kulkarni (Associate Professor), his student Dr. Alok D. Singh, and Dr. Shailesh Khade (Veterinarian) from Shobhaben Pratapbhai Patel School of Pharmacy & Technology Management (SPPSPTM), SVKM’s NMIMS, Mumbai for their support in helping with animal care and handling. We thank Mr. Shailesh Gala from Ms. Visha World (Lamington Rd, Grant Road East, Mumbai) for providing electronic parts for ERG instrumentation. We thank Dr. Kishore Dave (Shrushrusha Hospital, Dadar, Mumbai) for guidance on electrophysiological measurements. We also thank Mr Chandrashekhar Kulkarni, Associate Professor, Thadomal Shahani Engineering College Mumbai support in electrophysiological setup. for However, the article’s content, views, and conclusions contained herein are those of the primary and corresponding authors and are solely their responsibility.

## Author contributions

Dr. Anuradha V Pai performed the experiments and processed the data. Dr. Jayesh Bellare and Dr. Yogesh A. Kulkarni helped design the experiments. Dr. Jayesh Bellare contributed to data analysis and interpretation. Both Dr. Anuradha V Pai and Dr. Jayesh Bellare wrote the manuscript.

## Additional information

Extended data is available on request from the author.

## Supplementary

### Materials and Methods

#### Animal experiment

Procedures used in animal experiments were approved by Institutional Animal Ethics Committee of NMIMS [approval number CPCSEA/IAEC/P-16/2019] and conducted following the recommendation of the Association for Research in Vision and Ophthalmology Statement for the Use of Animals in Ophthalmology and Vision Research. Wistar rats (both gender, 2-2.5 months of age and 200-225 gm body weight: n = 6) were housed in a room with 12 hr light and dark cycle. Study animals received food and water ad libitum.

#### Animal preparation

Animals were be dark-adapted overnight. And all experimental setups were performed under dim red illumination for rod ERG and in −0.5 to 0.6 log cd/m^2^ (25-31 lux) and 0.5 log cd/m^2^ (316-318 lux) illumination for mesopic ERG and 1 log cd/m^2^ (1000 lux) for photopic ERG respectively. Rats were anesthetized with an intra-peritoneal ketamine (75-90 mg/kg) and xylazine (10 mg/kg) cocktail. Anesthesia with ketamine-xylazine was found to set within 5-10 mins for about 45-60 mins. In case required for prolonged study, anesthesia was maintained by repeat injection of 50% original dose of ketamine-xylazine every 40 mins. Pupils were dilated to 5 mm diameter with topical phenylephrine HCl (2.5%) and tropicamide (0.4%).

#### Procedure for intra-vitreal injection of drugs

Animals were kept under constant anaesthesia wrapped with a warm blanket to maintain its body temperature at 37 ± 2ᵒC (measured with the aid of a rectal thermometer). Two percent lidocaine was topically applied as local anaesthetic prior to intravitreal injections. Drugs (cis-PDA 100 mM / NaV modulators) were dissolved in 4 µl vehicle (normal saline pH 7.4) volume and was injected intravitreal using a specially designed pre-fillable cartridge with attached 30G needle which was loaded into a standard insulin pen. Our delivery device ensured precise error free injection volume and could be handled by single person. One eye was used to test while the other eye served as control (Xu et al., 2003). Concentration of lidocaine injected intravitreally was 85.35 mM, of lamotrigine was 600 µM and veratridine was 3.3 µM. Antibiotic (ciprofloxacin 0.3%w/v) was applied at the site of needle insertion with the help of earbud / cotton.

#### Recording system, light source, and calibrations

Rats were placed on a table (provided by SVKM’s NMIMS SPPSPTM, Mumbai) specially designed for acquiring ERG in order to keep head in a steady position and in order to reduce noise originating from respiratory and other movements and hence help reliable recording of ERG signals. Recording electrode was be made up of circular gold ring (4 mm diameter) corresponding to pupil of adult rat eye 4-5 mm diameter when dilated) and was be placed directly over the rat cornea. Stainless steel needle electrodes were used as reference as well as ground. Reference electrode was needle inserted into head skin of rat while the ground electrode was needle inserted into rat tail / muscle. Electrode contact was maintained using artificial tears (0.5% w/v carboxy methyl cellulose).

#### Method employed for eliciting ERG / stimulus condition

Protocol for all the ERG experiments were uniform for all groups of animals. ERG was recorded by means of Power Lab Data Acquisition system 2/25 (AD Instruments, New South Wales, Australia) and ERGs were acquired using LabChart version 7.3.8 (AD Instruments) for Windows, with a LabChart Pro license. A digital 50-60 Hz notch was applied to eliminate line noise. All the electrodes were connected to a differential amplifier (gain of 10,000; sampling rate set at 10 KHz; band pass filter 0.5 – 2 KHz (bridge amplifier)). ERGs were recorded against white flash of 509-515 nm wavelength and light calibrations were performed using a photometer (KUSAM-MECO; KM-LUX-99).

Dark-adapted rat ERG was recorded against single 10 msec (2 Hz) flash of white saturating full-field light stimulus (ranging from 1-1000 lux / −2 to 1 log cds / m^2^). The intervals between flashes were adjusted so that the response returned to baseline before another stimulus was presented. Animals were returned to their cage and constantly monitored to rule out eye infection / endophthalmitis due to intravitreal injection procedure. For measuring light-adapted ERG, a light source (Phillips LED) was mounted at pre-determined position above rat head to provide uniform illumination which was measured using digital photometer (KUSAM-MECO; KM-LUX-99).

**Fig. 1.**
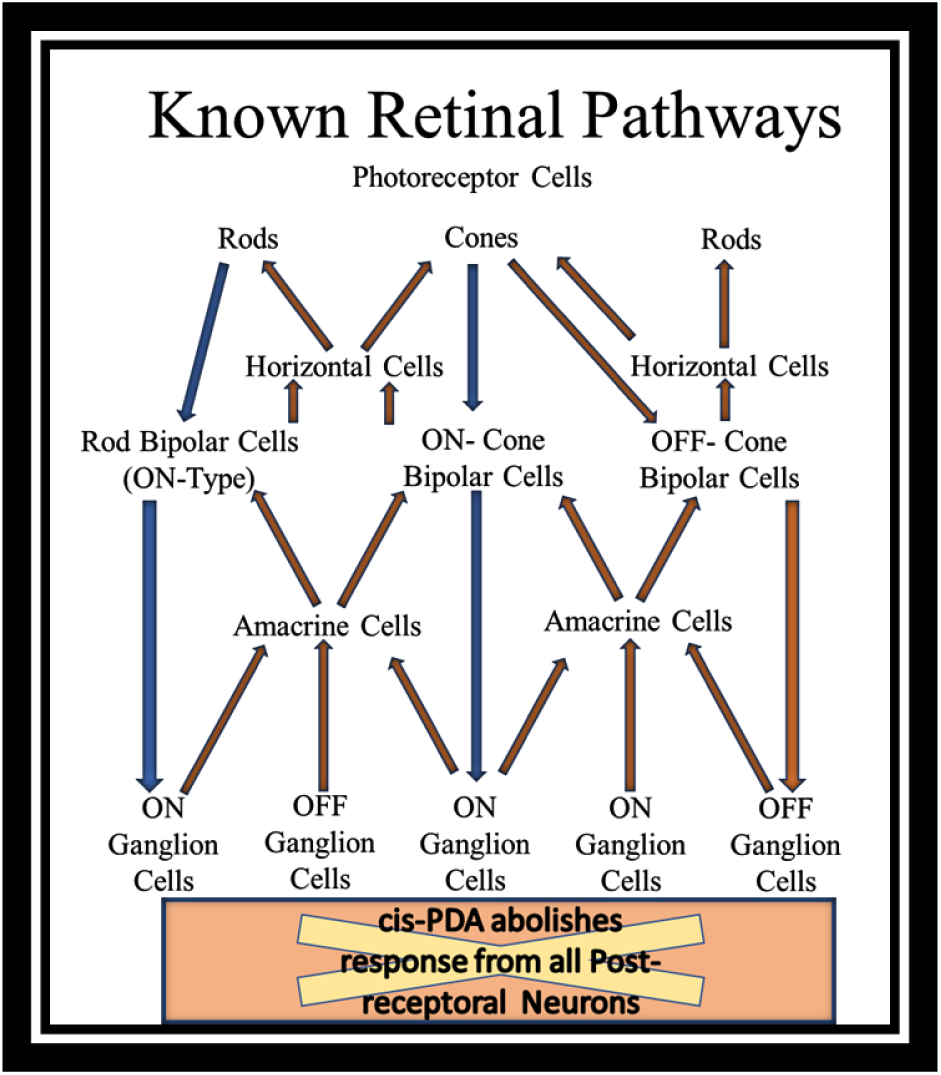
A simple retinal circuit and effect of cis-PDA. According to Xu et al, cis-PDA suppresses signal transmission from photoreceptors to OFF-cone bipolar cells (negative polarity response) and horizontal cells (feedback response of negative polarity) as well as between bipolar cells and third-order neurons (any feedback response of negative polarity) (all pink shaded response).

**Fig. 2.**
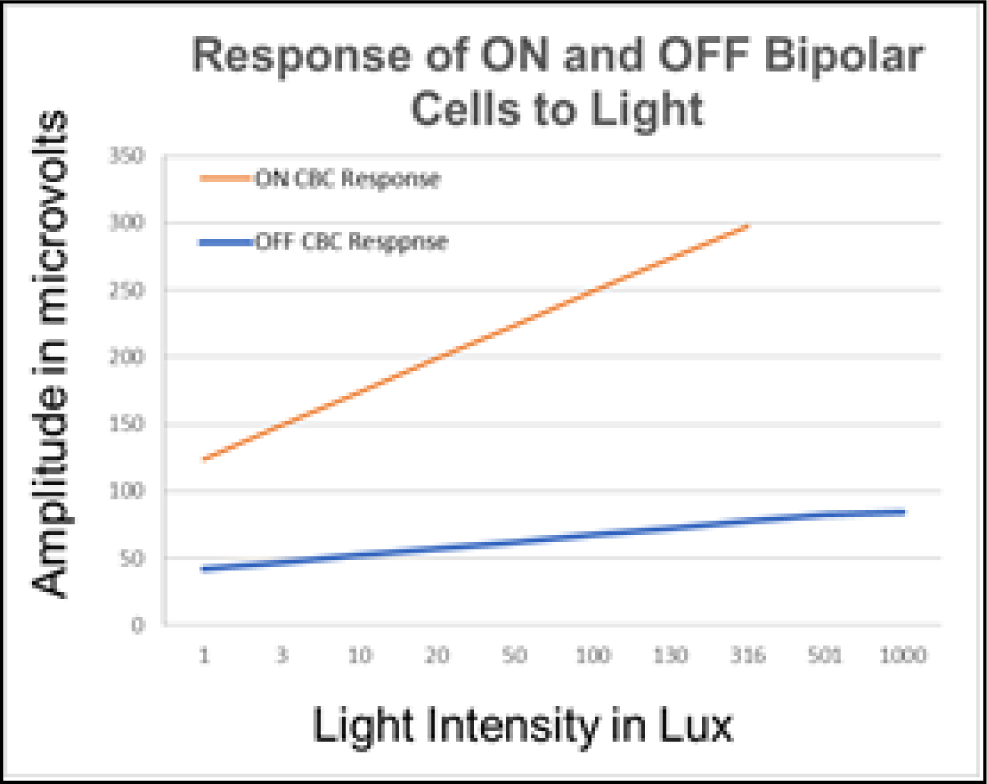
Response values of ON and OFF CBC to light stimulus of increasing intensity from 1-1000 lux.

**Table 1.** Different percentages of OFF and ON CBC image values combined (OFF CBC response added “+” or subtracted “-” from ON CBC response) for Image J Analysis [input data].

| ON →<br>OFF ↓ | 190 | 180 | 170 | 160 | 150 | 140 | 130 | 120 | 110 | 95 | 90 | 85 | 80 | 70 | 60 | 50 | 40 | 30 | 20 |
| --- | --- | --- | --- | --- | --- | --- | --- | --- | --- | --- | --- | --- | --- | --- | --- | --- | --- | --- | --- |
| -90 | +/- |  |  |  |  |  |  |  |  |  |  |  |  |  |  |  |  |  |  |
| -80 |  | +/- |  |  |  |  |  |  |  |  |  |  |  |  |  |  |  |  |  |
| -70 |  |  | +/- |  |  |  |  |  |  |  |  |  |  |  |  |  |  |  |  |
| -60 |  |  |  | +/- |  |  |  |  |  |  |  |  |  |  |  |  |  |  |  |
| -50 |  |  |  |  | +/- |  |  |  |  |  |  |  |  |  |  |  |  |  |  |
| -40 |  |  |  |  |  | +/- |  |  |  |  |  |  |  |  |  |  |  |  |  |
| -30 |  |  |  |  |  |  | +/- |  |  |  |  |  |  |  |  |  |  |  |  |
| -20 |  |  |  |  |  |  |  | +/- |  |  |  |  |  |  |  |  |  |  |  |
| -10 |  |  |  |  |  |  |  |  | +/- |  |  |  |  |  |  |  |  |  |  |
| 5 |  |  |  |  |  |  |  |  |  | +/- | +/- | +/- | +/- | +/- | +/- | +/- |  |  |  |
| 10 |  |  |  |  |  |  |  |  |  | +/- | +/- | +/- | +/- | +/- | +/- | +/- |  |  |  |
| 15 |  |  |  |  |  |  |  |  |  | +/- | +/- | +/- | +/- | +/- | +/- | +/- |  |  |  |
| 20 |  |  |  |  |  |  |  |  |  | +/- | +/- | +/- | +/- | +/- | +/- | +/- |  |  |  |
| 30 |  |  |  |  |  |  |  |  |  | +/- | +/- | +/- | +/- | +/- | +/- | +/- |  |  |  |
| 40 |  |  |  |  |  |  |  |  |  | +/- | +/- | +/- | +/- | +/- | +/- | +/- |  |  |  |
| 50 |  |  |  |  |  |  |  |  |  | +/- | +/- | +/- | +/- | +/- | +/- | +/- |  |  |  |
| 60 |  |  |  |  |  |  |  |  |  |  |  |  |  |  |  |  | +/- |  |  |
| 70 |  |  |  |  |  |  |  |  |  |  |  |  |  |  |  |  |  | +/- |  |
| 80 |  |  |  |  |  |  |  |  |  |  |  |  |  |  |  |  |  |  | +/- |

